# neoepiscope improves neoepitope prediction with multi-variant phasing

**DOI:** 10.1101/418129

**Authors:** Mary A. Wood, Austin Nguyen, Adam Struck, Kyle Ellrott, Abhinav Nellore, Reid F. Thompson

**Affiliations:** Computational Biology Program, Oregon Health & Science University; Portland VA Research Foundation; Department of Biomedical Engineering, Oregon Health & Science University; Department of Surgery, Oregon Health & Science University; Department of Radiation Medicine, Oregon Health & Science University; Department of Medical Informatics & Clinical Epidemiology, Oregon Health & Science University; Division of Hospital and Specialty Medicine, VA Portland Healthcare System

**Author notes:** co-corresponding authors [, ].

## Abstract

The vast majority of tools for neoepitope prediction from DNA sequencing of complementary tumor and normal patient samples do not consider germline context or the potential for co-occurrence of two or more somatic variants on the same mRNA transcript. Without consideration of these phenomena, existing approaches are likely to produce both false positive and false negative results, resulting in an inaccurate and incomplete picture of the cancer neoepitope landscape. We developed `neoepiscope` chiefly to address this issue for single nucleotide variants (SNVs) and insertions/deletions (indels), and herein illustrate how germline and somatic variant phasing affects neoepitope prediction across multiple datasets. We estimate that up to ∼5% of neoepitopes arising from SNVs and indels may require variant phasing for their accurate assessment. `neoepiscope` is performant, flexible, and supports several major histocompatibility complex binding affinity prediction tools. We have released `neoepiscope` as open-source software (MIT license, https://github.com/pdxgx/neoepiscope) for broad use.

**KEY POINTS:**

- Germline context and somatic variant phasing are important for neoepitope prediction
- Many popular neoepitope prediction tools have issues of performance and reproducibility
- We describe and provide performant software for accurate neoepitope prediction from DNA-seq data

## INTRODUCTION

While mutations may promote oncogenesis, cancer-specific variants and the corresponding novel peptides they may produce (“neoepitopes”) appear central to the generation of adaptive anti-tumor immune response [1]. The advent of immunotherapy as a promising form of cancer treatment has been accompanied by a parallel effort to explore tumor neoepitopes as potential mechanisms and drivers of therapeutic response. This has prompted the development of numerous neoepitope prediction pipelines. Initial bioinformatics approaches used tumor-specific sequencing data to focus on missense single nucleotide variants (SNVs) as the predominant source of neoepitopes (e.g. Epi-Seq [2]). However, as ∼15% of neoepitopes are estimated to result from other types of mutations [3], additional tools were developed to predict neoepitopes from gene fusions (e.g. INTEGRATE-neo [4]), non-stop mutations (e.g. TSNAD [5]), and insertions and deletions (indels, from e.g. pVACseq [6], MuPeXI [7]), which may be of particular significance for anticipating cancer immunotherapy response [8]. Many of these tools enable comparable predictive capabilities, but each approach has its own unique set of features and limitations (See Figure 1).

**Figure 1:**
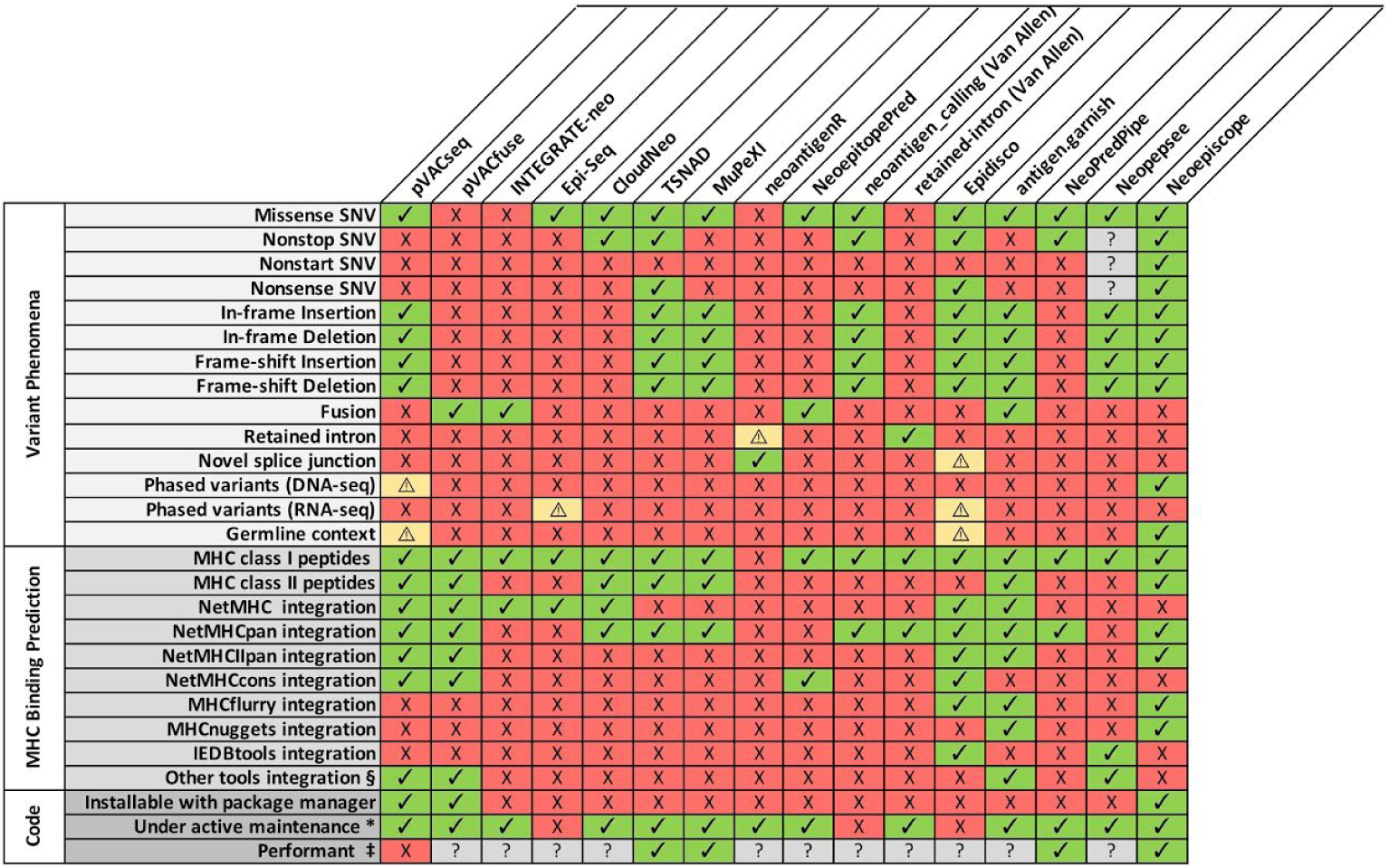
A feature comparison of pVACseq and pVACfuse [6], INTEGRATE-neo [4], Epi-Seq [2], CloudNeo [58], TSNAD [5], MuPeXI [7], neoantigenR [59], NeoepitopePred [60], neoantigen_calling_pipeline (Van Allen laboratory) [27], retained-intron-neoantigen-pipeline (Van Allen laboratory) [52], Epidisco [61], antigen.garnish [62], NeoPredPipe [63], Neopepsee [64], and `neoepiscope`. Each column corresponds to a software tool, with software features listed by row. A green check mark indicates that the tool possesses or processes the indicated feature, while a red “X” indicates that the tool does not possess or process the indicated feature. A yellow warning symbol 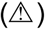 indicates that a tool incompletely supports the corresponding feature. Specifically, neoantigenR can only identify retained introns when using GTF annotations that contain labeled introns. Epidisco is limited to RNA-seq data and cannot account for downstream effects of a variant, though it can identify some events of alternative splicing, germline variation, and nearby somatic variation so long as they occur nearby a declared variant. Epi-Seq uses phasing information to determine the genotypes of variants and to identify other variants nearby variants that produce neoepitopes, but like Epidisco it does not take into account the downstream effects of a variant. pVACseq is able to perform variant phasing for substitutions including proximal germline variants using GATK ReadBackedPhasing [65]. Gray question marks (“?”) denote unknown or unassessed values. * A tool was considered to be under activate maintenance if a new release or GitHub commit had occurred within 6 months prior to submission of this manuscript. § Other MHC binding prediction tools used include NNalign, PickPocket, SMM, SMMPMBEC, and SMM align for pVACseq and pVACfuse; NetMHCII for antigen.garnish, and NetCTLpan for Neopepsee. ‡ A tool was considered to be performant if neoepitope prediction averaged less than 10 minutes per sample in our benchmarking (see Methods).

However, a limitation of most neoepitope prediction tools is that they consider individual somatic variants in the sequence context of a single reference genome, neglecting interactions between somatic and potentially neighboring germline variants given the widespread genetic variability among individuals [9]. Indeed, somatic and germline mutations may co-occur even within the 3 nucleotide span of a single codon [10]. Such immediate co-occurrences are not uncommon, affecting 17% of cancer patients on average, and can significantly alter variant effects including the amino acid sequence of any predicted neoepitope. Somatic mutations may also cluster or co-occur within a short span of each other. For instance, several hundred somatic mutations were found to co-occur within a <10bp intervariant distance among whole genome sequencing data from two breast cancer patients [11]. Furthermore, immediate co-occurrences of variants are not a prerequisite for phasing to be relevant to neoepitope prediction. For example, frameshifting indels upstream of a SNV can alter the peptide sequence in the vicinity of the SNV, as well as the consequences of the SNV itself.

The vast majority of neoepitope prediction tools only consider single somatic variants in isolation from each other, neglecting the potential for these compound effects. We developed `neoepiscope` chiefly to incorporate germline context and address variant phasing for SNVs and indels, and herein illustrate how these features affect neoepitope prediction across multiple patient datasets.

## METHODS

### Software

The `neoepiscope` neoepitope prediction pipeline is depicted in Figure 2. It is highly flexible and intentionally agnostic to the user’s preferred tools for alignment, human leukocyte antigen (HLA) typing, variant calling, and MHC binding affinity prediction. The pipeline upstream of `neoepiscope` takes as input DNA-seq (exome or whole-genome) FASTQs for a tumor sample and a matched normal sample. These FASTQs are used to perform HLA genotyping, e.g. using OptiType [12]. The FASTQs are also aligned to a reference genome with a DNA-seq aligner such as BWA [13] or Bowtie 2 [14]. The resulting alignments are subsequently used to call somatic and, optionally, germline variants. Somatic variants are obtained by running any somatic variant caller or combination of callers on the tumor and normal sample’s alignments, acknowledging that consensus-based approaches give more accurate results than a single caller alone [15]. To obtain In most of our results, we used a combination of Pindel, MuSE, MuTect, RADIA, SomaticSniper, and VarScan 2 (see *Variant identification and phasing*). Germline variants are obtained by running a germline variant caller such as Genome Analysis Toolkit’s (GATK) HaplotypeCaller [16] or VarScan 2 [17] on the normal sample’s alignments. Next, somatic variants are phased, optionally with germline variants, by a variant phasing tool such as HapCUT2 [18] (note that `neoepiscope` was designed to be run with HapCUT2-formatted input, but can also be run with a phased VCF produced by GATK’s ReadBackedPhasing or an unphased VCF). The resulting phased variants (“phase sets”) are input to `neoepiscope` together with a gene transfer format (GTF) file encoding known transcript variants (e.g., GENCODE).

**Figure 2:**
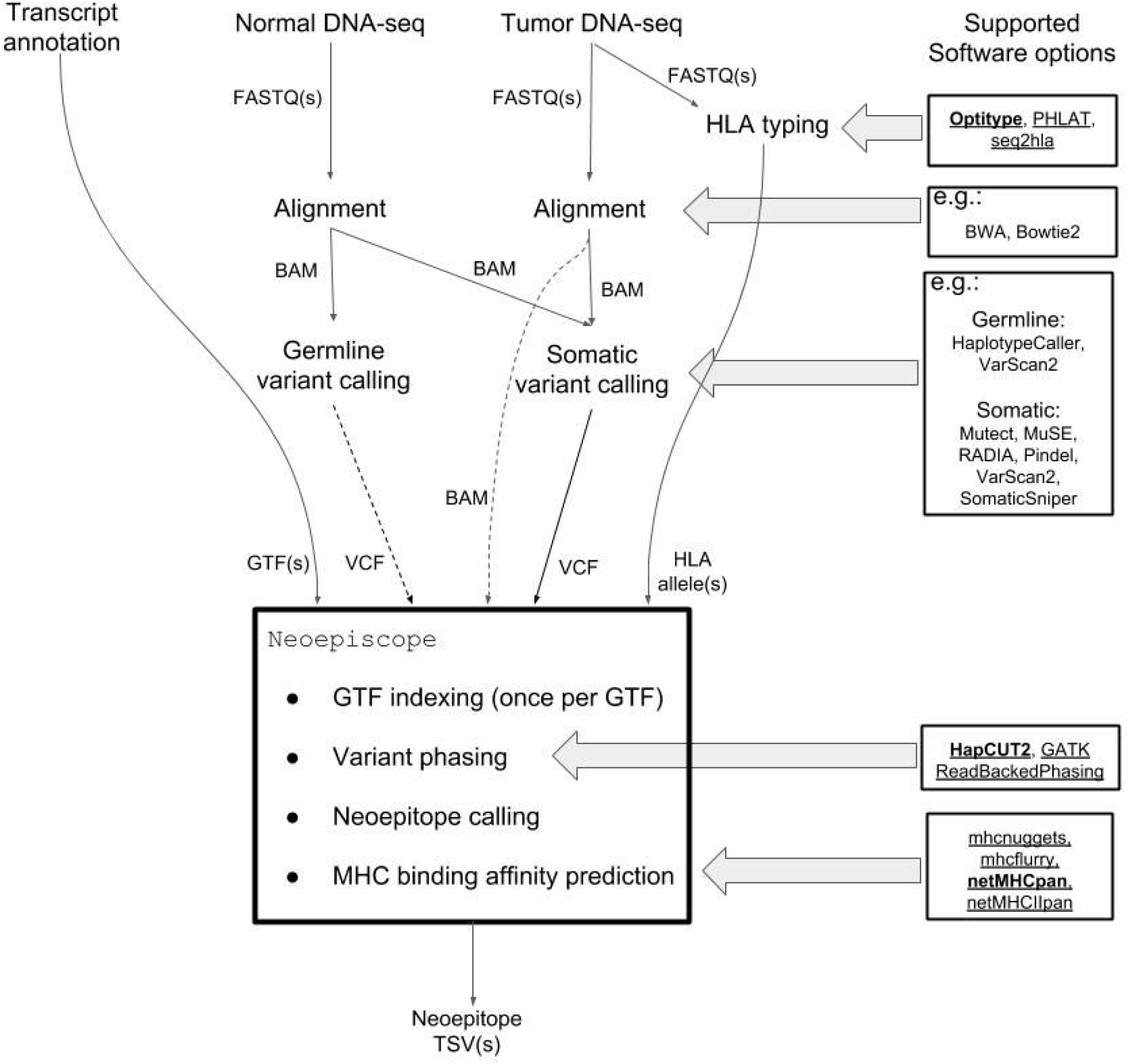
Neoepitope prediction pipeline diagram describing canonical `neoepiscope` workflow. Global inputs are shown at the top of the figure, with connecting arrows demonstrating interim inputs and outputs between processing steps. Direct inputs to and outputs from `neoepiscope` are shown directly entering or leaving the outlined box listing `neoepiscope` functionality. Multiple potential software options are shown at right for each relevant processing step as indicated by horizontal arrows (tools that are directly compatible with `neoepiscope` are underlined, with those in bold implemented as default).

The principal contribution of this manuscript is the neoepitope calling component of `neoepiscope`. Before it is run on the phase sets associated with a given tumor-normal sample pair, `neoepiscope` indexes the GTF into a set of pickled dictionaries that link exon, start codon, and stop codon blocks from transcripts to the genomic intervals they cover. These dictionaries can be reused with every call to `neoepiscope` to reduce runtime. Prior to neoepitope prediction, unphased somatic variants from the somatic VCF are added to HapCUT2 output as their own haplotype blocks using `neoepiscope`’s prep mode. For each haplotype block, `neoepiscope` determines if any mutation in the block overlaps any transcript(s) in the annotation, and then applies each mutation to the transcript. (Transcript sequences are obtained by concatenating relevant genomic sequence from an appropriate Bowtie [19] index). By default, all relevant somatic and germline variants in a haplotype block are applied, but users may choose to exclude either germline or somatic variants, or to exclude variant phasing entirely. Each transcript is translated *in silico*, proceeding from the annotated start codon (or the first annotated codon if no start is present), but this process may be modified via command line options to perform alternative start codon predictions. The resulting protein is kmerized in the vicinity of amino acid changes, and epitopes that do not match any peptide sequence in the normal protein are returned as potential neoepitopes. We performed extensive unit testing on our software to ensure that both common and rare cases of somatic variants are handled accurately. Patient-specific HLA alleles and a choice of MHC binding prediction tools can be specified so that optional binding prediction can be performed for each potential neoepitope. As indicated in Figure 2, `neoepiscope` itself calls the user-specified MHC binding affinity prediction software to obtain HLA allele-associated binding affinities. By default, it installs MHCflurry [20] and MHCnuggets [21]. The user may also independently install NetMHCpan [20] or NetMHCIIpan [22] for use with `neoepiscope`.

### Variant identification and phasing

We assembled a cohort of 375 tumor samples from 347 different cancer patients from publicly available data including 285 melanoma patients (313 tumor samples) [23–30], 34 non-small cell lung cancer patients (28 tumor samples) [31], and 28 colon, endometrial, and thyroid cancer patients (29 tumor samples) [32] (see Supplementary Table 1). Whole exome sequencing (WES) reads for each sample were aligned to GRCh37d5 using the Sanger cgpmap workflow [33], which uses bwa-mem (v0.7.15-1140) [34] and biobambam2 (v2.0.69) [35] to generate genome-coordinate sorted alignments with duplicates marked and Genome Analysis Toolkit (GATK, v3.6) to realign around indels and perform base recalibration for paired tumor and normal sequence read data. Somatic variants were called and filtered using MuSE (v0.9.9.5) [36], MuTect (v1.1.5) [37], Pindel (v0.2.5b8) [38], RADIA (v1.1.5) [39], SomaticSniper (v1.0.5.0) [40], and VarScan 2 (v2.3.9) [17] according to the mc3 variant calling protocol [41], a consensus-based gold standard approach adapted from crowdsourced benchmarking data (i.e. DREAM challenge) [15]. To make somatic variants reported across tools comparable, vt (v0.5772-60f436c3) [42] was used to normalize variants, decompose biallelic/block variants, sort variants, and produce a final, unique list of variants for each caller. Somatic variants that were reported by more than one tool, or that were reported by Pindel and overlapped at least one call from another tool, were retained for further analysis. Germline variants were called using GATK’s HaplotypeCaller (v3.7-0-gcfedb67) [16], and we used VariantFiltration with cluster size 3 and cluster window size 15 to flag and subsequently eliminate variants with total coverage of less than 10.0 reads, quality by read depth of less than 2.0, and phred quality of less than 100.0. For consistency, germline variants were also processed with vt as described above. The Mbp of genome covered was determined using bedtools genomecov (v2.26.0) [43] where any base covered by a depth of at least 3 reads was considered covered, as this is the minimum read depth required for variant detection by SomaticSniper and VarScan 2. The coverage-adjusted mutational burden was calculated as the number of somatic mutations per Mbp of genome covered. We employed HapCUT2 for patient-specific haplotype phasing. To do this, germline and somatic variants were combined into a single VCF using `neoepiscope`’s merge functionality. HapCUT2’s extractHAIRS software was run with the merged VCF and the tumor alignment file, allowing for extraction of reads spanning indels, to produce the fragment file used with HapCUT2 to predict haplotypes. Neoepitopes were predicted for this cohort using the default settings of `neoepiscope`, either including germline variation and variant phasing (“phased mode”) or using unphased somatic variants only (“unphased mode”) in order to determine how phasing of variants might impact neoepitope predictions. The proportion of false positive neoepitope predictions when not accounting for variant phasing was calculated as the number of neoepitopes predicted by the unphased mode but not the phased mode, divided by the total number of neoepitopes predicted by the unphased mode. The proportion of false negative neoepitope predictions was calculated as the number of neoepitopes predicted by the phased mode but not the unphased mode, divided by the total number of neoepitopes predicted by the unphased mode.

For 90 patients [24–27], complementary RNA sequencing (RNA-seq) data was available in addition to WES data (see Supplementary Table 1). We aligned these RNA-seq reads to the hg19 genome using STAR (v2.6.1c) [44] and searched for evidence of phasing of predicted somatic variants with germline or other somatic variants within the span of a single read pair to confirm predicted phasing by HapCUT2 and identify instances of phasing not detected by HapCUT2. If two variants were both present in at least one paired read, they were considered to be phased in RNA. If one of two variants predicted to be phased was missing from a read, this was considered evidence for lack of phasing only if 1) the variant present in the read was somatic and 2) there were reads from that transcript that contained at least one somatic mutation, as contaminating normal tissue could explain the presence of a germline variant alone. Scripts to identify phasing of variants within RNA-seq reads are available at https://github.com/pdxgx/neoepiscope-paper under the MIT license.

### Benchmarking

We compared the performance of `neoepiscope` with pVACseq (pvactools v1.0.7) [6] and MuPeXI (v1.2.0) [7] on five melanoma patients for whom paired tumor and normal DNA-seq data were available [45]. Because MuPeXI was designed to work most readily with the GRCh38 genome build, we benchmarked all software using this build, which required several modifications of the approach described in “*Variant identification and phasing*” above. Paired tumor and normal WES reads were each aligned to the GRCh38 genome using bwa mem (v0.7.9a-r786) [34], and GATK (v4.0.0.0) [16] was used to mark duplicates and perform base recalibration. Somatic variants were called using MuTect2 (GATK v3.5-0-g36282e4) [37], and germline variants were called using GATK HaplotypeCaller (v3.5-0-g36282e4), with VariantFiltration performed as described in “*Variant identification and phasing*”. HLA types were predicted from tumor WES reads using Optitype (v1.3.1) [12]. Patient-specific haplotypes were predicted using HapCUT2 [18] as described in “*Variant identification and phasing*”, both with and without the inclusion of germline variation. For use with pVACseq and TSNAD, somatic VCFs were annotated with Variant Effect Predictor (v91.3) [46] and variants were phased using GATK’s ReadBackedPhasing according to pVACseq recommendations. Neoepitope prediction with pVACseq, MuPeXI, NeoPredPipe, TSNAD, and `neoepiscope` was performed with the most updated versions/commits of the software available on March 4, 2019. We ran pVACseq with phasing of somatic and germline variants, and `neoepiscope` in comprehensive mode. Across all tools, HLA binding predictions were performed with netMHCpan (v4.0 for MuPeXI/`neoepiscope`, v2.8 for pVAC-Seq, which uses IEDB Tools’ netMHCpan integration) [47] for consistency with MuPeXI’s requirements using alleles HLA-A*01:01, HLA-B*08:01, and HLA-C*07:01 (shared across four of the five patients). A uniform peptide length of 9 amino acids was used for prediction in all cases, with a maximum binding score cutoff of 5×10^4^ nM. Performance time and predicted neoepitope sequences were compared across tools (Figures 5 and 6), using four Intel Xeon E5-2697 v2 processors (when multithreading was possible), each with twelve 2.70 GHz processing cores. Scripts to reproduce the benchmarking described here are available at https://github.com/pdxgx/neoepiscope-paper under the MIT license. Predicted neoepitope sequences were compared across tools, and neoepitopes missed by any individual tool(s) were investigated to evaluate accuracy. Neoepitope sequences deriving from phased variants were considered accurate (true positives) if their phasing was confirmed by both HapCUT2 and ReadBackedPhasing; their accuracy was considered unknown wherever HapCUT2 and ReadBackedPhasing disagreed. If a neoepitope spanning the positions of two accurately phased variants was derived from only one half of the pair, that neoepitope sequence was considered to be a false positive, often accompanied by a false negative corresponding to the neoepitope sequence incorporating the pair of phased variants. The accuracy of neoepitope sequences deriving from putative RNA-editing events or mutations in unlocalized contigs was considered unknown.

**Figure 3:**
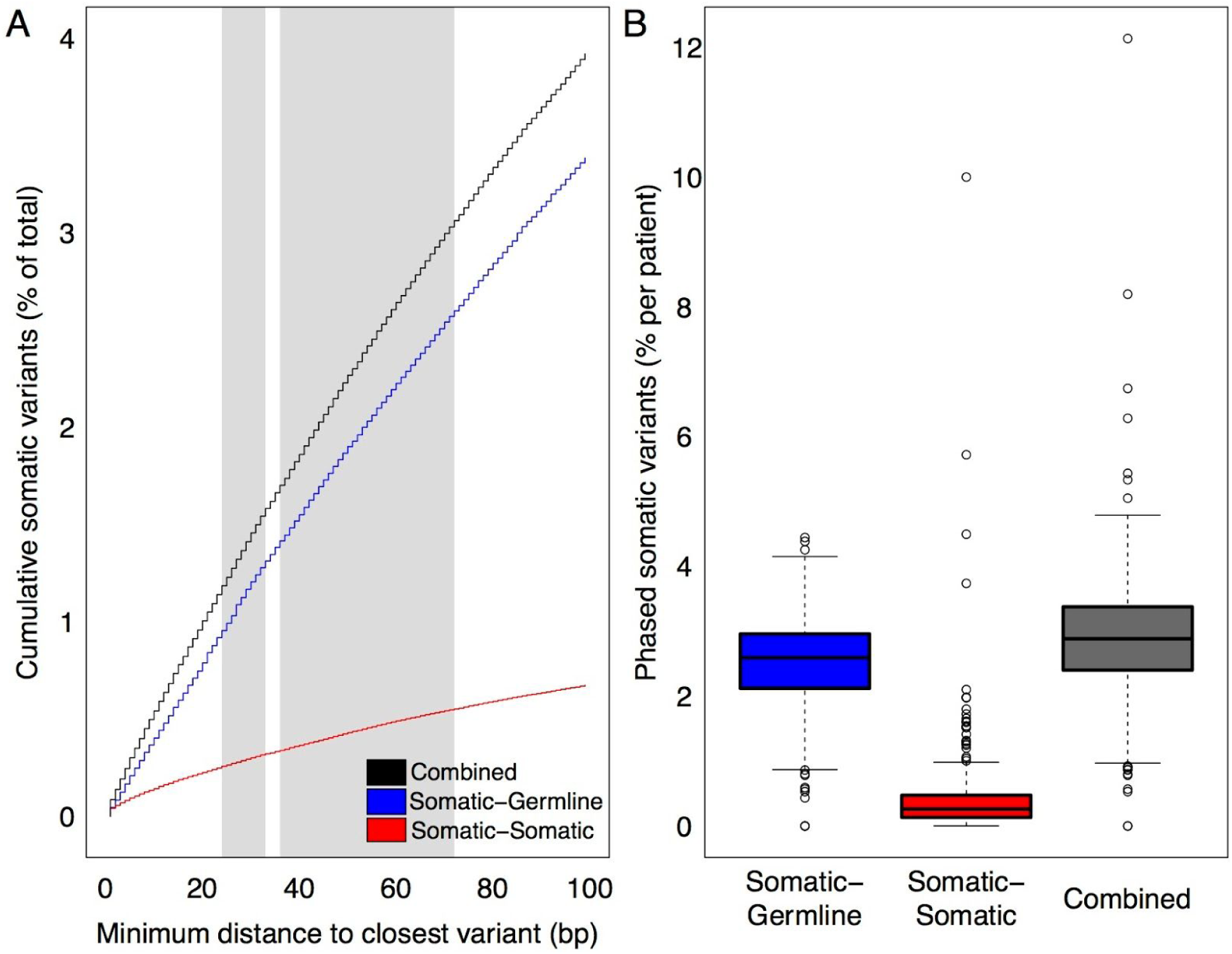
Variant co-occurrence among 285 melanoma patients, 34 NSCLC patients, and 28 colon, endometrial, and thyroid cancer patients. A) The cumulative average percentage (y-axis) of somatic variants across all tumors that co-occur with germline variants (blue), other somatic variants (red), or either type of variant (black) is shown as a function of increasing nucleotide span (x-axis). Canonical MHC Class I and Class II epitope size ranges are shaded in light gray (24-33bp and 36-72bp, respectively). B) Box plots demonstrating per-patient percentage of somatic variants (y-axis) across all tumors that co-occur with germline variants (blue), other somatic variants (red), or either variant type (dark gray).

**Figure 4:**
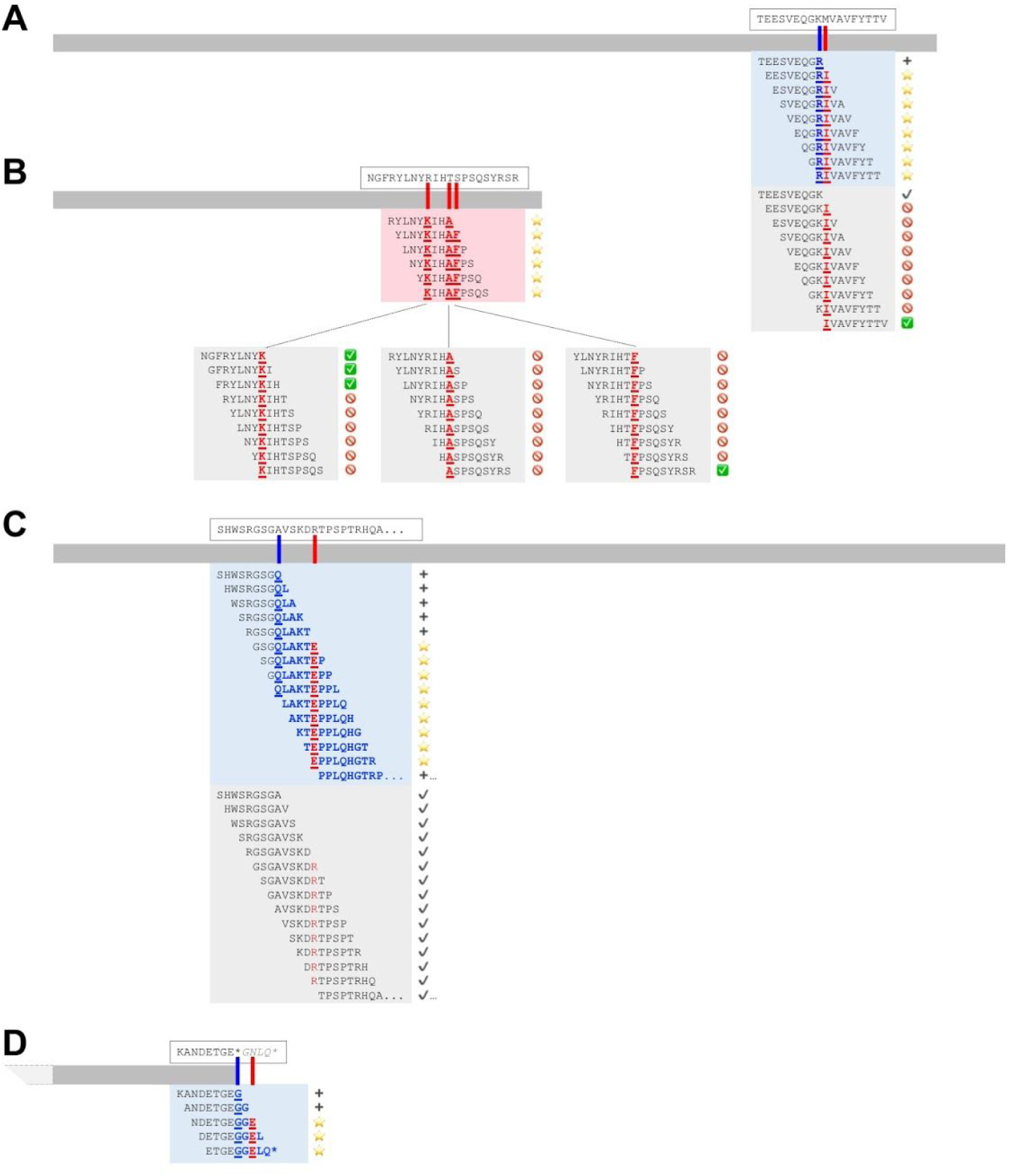
Neoepitope-level consequences of phased variants. Four coding sequences are shown to relative scale (gray horizontal bars), with corresponding approximate variant locations shown in red or blue tick marks for somatic and germline variants, respectively. Amino acid sequences corresponding to the reference coding sequence are shown in dashed boxes above their corresponding transcripts, with “…” indicating additional unspecified sequence, a ^“*”^ denoting a stop codon, and italicized characters indicating untranslated sequence. Colored highlighted boxes contain 9mer neoepitopes and their immediate context as predicted from phased variants, with shading corresponding to somatic variants alone (red) or including germline context (blue). Gray highlighted boxes contain 9mer neoepitopes and their immediate context as predicted from somatic variants in isolation, without consideration of phasing or germline context. Amino acid sequences that are directly affected by germline or somatic variants are displayed in bold underlined blue or red, respectively, while downstream variant consequences are shown without underline, and silent variant effects are neither bolded nor underlined. Corresponding symbols are shown at the right of each peptide and demonstrate the anticipated consequence of incorporating variant phasing and germline context, with yellow stars denoting novel neoepitopes, red NOT symbols denoting incorrectly predicted neoepitopes, green checkboxes denoting correctly predicted neoepitopes, black plus signs denoting novel proteomic context, and black check boxes denoting consistent proteomic context. Co-occurring variant phenomena correspond to real patient data as noted in the text and are depicted as follows: A) germline SNV + somatic SNV, B) multiple somatic SNVs, C) germline frameshift deletion + downstream somatic SNV, and D) germline non-stop variant + downstream somatic SNV.

**Figure 5:**
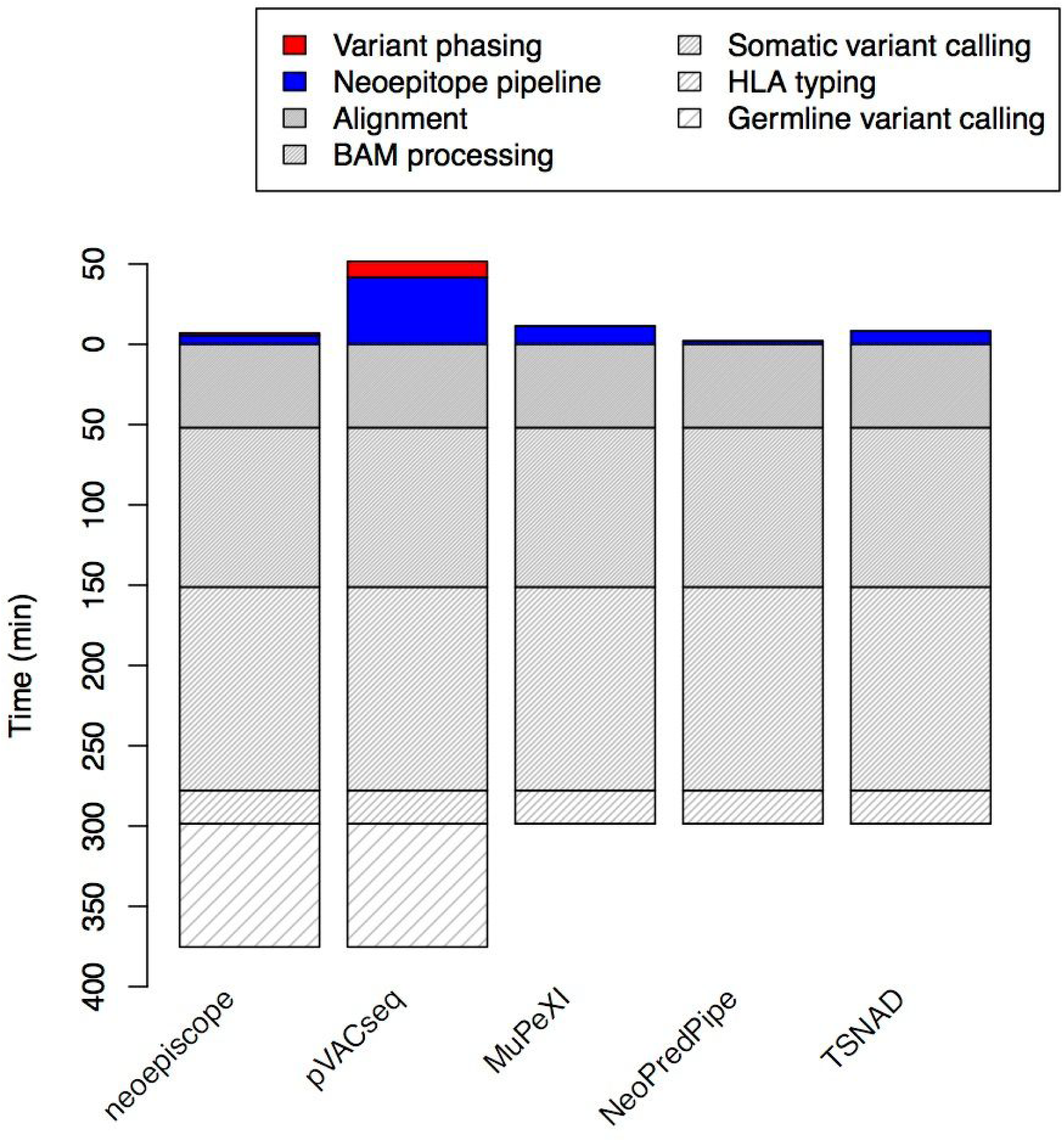
Performance benchmarking of `neoepiscope`, pVACseq, MuPeXI, NeoPredPipe, and TSNAD. The run time (y axis) of `neoepiscope` including germline variation was compared with those of pVACseq, MuPeXI, NeoPredPipe, and TSNAD at a pipeline level. Time zero indicates the beginning of the neoepitope calling process from somatic (and germline, if relevant) variants, such that steps in the pipeline prior to neoepitope prediction (e.g. sequence read alignment, somatic variant calling, etc.) are displayed below time zero, and steps during or required for neoepitope calling are displayed above time zero. Only `neoepiscope` and pVACseq pipelines required variant phasing. Variant Effect Predictor (VEP) runtimes were included in the pVACseq and TSNAD neoepitope prediction steps (VEP is a prerequisite for pVACseq and TSNAD).

**Figure 6:**
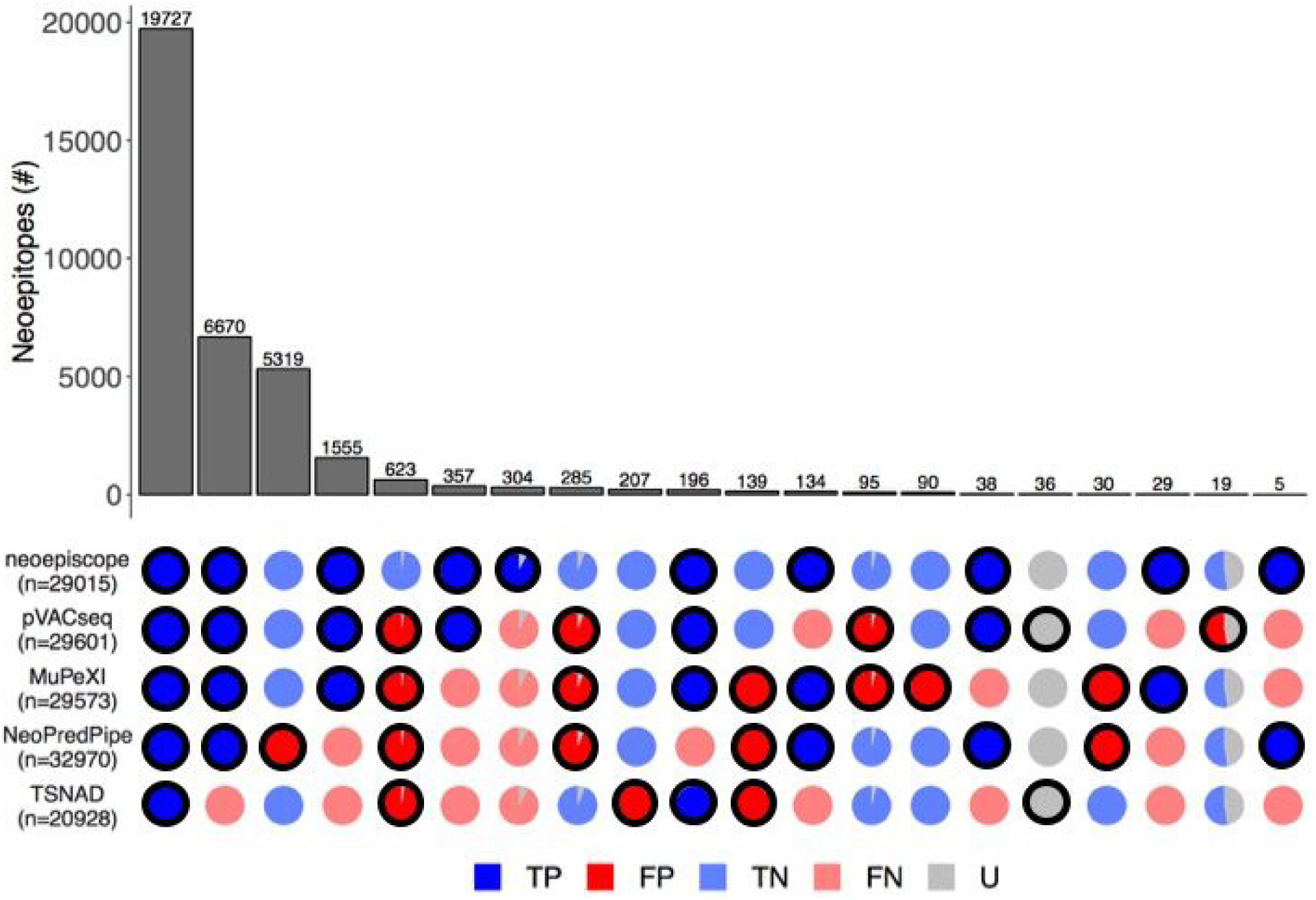
Detailed comparison of the complete set of neoepitope predictions from `neoepiscope`, MuPeXI, pVACseq, NeoPredPipe, and TSNAD. Patterns of agreement or disagreement among groups of neoepitopes predicted by different combinations of tools across 5 melanoma patients are shown along each column (e.g. the first column corresponds to neoepitopes predicted by all tools). Each row indicates the neoepitope predictions associated with the indicated tool, with the total number of neoepitopes predicted by each tool shown as *n*. The number of neoepitopes in each column (bar in upper pane) corresponds to the size of the subset predicted by the indicated combination of tools (outlined circles in the bottom pane). The veracity of predictions corresponding to each group of neoepitopes are shown as pie charts, with colors corresponding to true positive (“TP”, dark blue), false positive (“FP”, dark red), true negative (“TN”, light blue), false negative (“FN”, light red), and uncertain (“U”, gray) predictions. Uncertain predictions are considered a result of one or more factors including origin from unassembled contig regions, the presence of RNA edits, or inconsistencies in variant phasing predictions between HapCUT2 and GATK’s ReadBackedPhasing.

### Estimation of variant co-occurrence rates

To estimate patient-specific variant co-occurrence, predicted haplotypes were used to determine when somatic variants were proximal to either a germline variant or another somatic variant within the span of an MHC-II epitope (72 bp/24 aa) in the cohort described in *“Variant identification and phasing”*. Because of variant decomposition during somatic variant calling, immediately adjacent somatic single nucleotide variants predicted to be in the same alleles were collapsed into a single multiple nucleotide variant. Nearby germline variants were counted both upstream and downstream of somatic variants, while somatic variants were only counted upstream of each other to avoid duplication. Proximity of frameshift variants and nonstop variants was considered based on strand specificity of any transcripts overlapping a somatic variant, such that only the effects of upstream frameshifts and nonstops were considered for downstream variants. Both the number of variants near a somatic variant and the minimum distance between a somatic variant and its nearby variants were recorded.

## RESULTS

### Somatic and germline variant co-occurrence

We analyzed the intervariant distance distribution among somatic and germline variants from 375 tumor samples, employing HapCUT2 [18] for patient-specific haplotype phasing (see Methods). We found that variant co-occurrence increases approximately linearly with increasing nucleotide span, with an overall average of 2.72% and 0.43% of somatic variants located within a 72bp inter-variant distance of another germline or somatic variant, respectively (Figure 3A). On a per-patient basis, as many as 3.25% and 5.32% of somatic variants co-occurred with at least one germline variant within the span of an MHC class I (≤33bp) and class II (≤72bp) epitope, respectively (Figure 3B). Additionally, as many 9.15% and 10.1% of somatic variants per patient co-occurred with at least one other somatic variant within the span of an MHC class I or class II epitope. For a subset of patients (n=90), we also analyzed RNA-seq data to assess phasing across exon-exon junctions (see Methods). Acknowledging that many variants may occur in genes that are not expressed in tumor samples, only 20.5% of phased somatic-germline variant pairs and only 38.9% of phased somatic-somatic variant pairs were covered by aligned RNA-seq reads and present in tumor-expressed transcripts. Among variant pairs phased across exon-exon junctions using DNA-seq data, an average of 80.9% of those covered by RNA-seq reads were able to be validated by corresponding RNA-seq data, while a small fraction of those covered (2.24%) demonstrated discordant phasing. RNA-seq also identified an additional 0.18% of phased variant pairs on average across exon-exon junctions, respectively, that were not identified using DNA-seq.

The percentage of phased somatic variants was not significantly different between disease types (p = 0.758 by ANOVA; see Supplementary Figure 1), but did correlate with tumor mutational burden (Pearson’s product-moment correlation of 0.35, p = 2.93×10^−11^; see Supplementary Figure 2). Based on these data, we predicted neoepitope sequences either with or without considering variant phasing, and found that neoepitope prediction without consideration of germline context or variant phasing produces a median of 4.25% false positive results and 2.41% false negative results (see Methods), resulting in both an inaccurate and incomplete picture of the cancer neoepitope landscape.

### Neoepitope prediction consequences of variant phasing

We identified specific examples of co-occurring variants from real patient data, as above, and show the `neoepiscope`-predicted consequences of both variant phasing and germline context for each of these examples in Figure 4. Co-occurrence of neighboring somatic and germline SNVs can produce neoepitopes with multiple simultaneous amino acid changes. For instance, germline (chr11:56230069 T->C) and somatic (chr11:56230065 C->T) variants in a melanoma patient (“Pat41” [27]) give rise to a collection of potential compound neoepitopes containing two adjoining amino acid changes (germline K270R and somatic M271I in ENST00000279791) (Figure 4A). Such hybrid (somatic+germline) co-occurrence within the span of an MHC Class I or Class II epitope accounts for an average of 2.72% of somatic variants overlapping transcripts (range: 0-5.32%).

Three neighboring somatic variants (chr21:31797833 C->T, chr21:31797825 T->C, and chr21:31797821 G->A) in another melanoma patient (“37” [29]) give rise, jointly, to three amino acid changes (R133K, T136A, and S137F in ENST00000390690) (Figure 4B). In such cases of neighboring SNVs, the anticipated proportion of compound neoepitopes is inversely and linearly related to the distance separating neighboring variants on both sides (Figure 4B). Co-occurrence of two or more somatic variants accounts for an average of 0.43% of somatic variants overlapping transcripts (range: 0-10.1%).

It is also possible for frameshift indels to be located within upstream coding sequences near one or more phased variants. For instance, a somatic SNV (chr10:76858941 C->T) in the context of an upstream germline frameshifting deletion (chr10:76858961 CT->C) in another melanoma patient (“Pt3” [25]) introduces multiple novel potential neoepitopes arising from ENST00000478873 (Figure 4C). Importantly, without germline context, this somatic SNV is predicted to be a silent mutation and would not otherwise be predicted to give rise to any neoepitopes. We analyzed the prevalence of frameshift indels neighboring downstream variants within a 94bp window (distance corresponding to an estimated ∼95% probability of encountering a novel out-of-frame stop codon). On a per patient basis, an average of 0.33% of somatic variants overlapping transcripts were phased with an upstream frameshifting indel (range: 0-1.16%).

Finally, stop codon mutations that enable an extension of the reading frame may expose one or more downstream somatic or germline variants. For instance, a germline SNV (chr3:17131402 T->G) in another melanoma patient (“Pt1” [25]) eliminates a stop codon in ENST00000418129, with an anticipated extension of translation of an additional 5 amino acids prior to encountering the next stop codon. However, the co-occurrence of a downstream somatic SNV (chr3:17131406 G->A) alters the anticipated peptide composition (Figure 4D). We analyzed the frequency of such non-stop variants co-occurring with one or more downstream variants within a 94bp window (distance corresponding to an estimated ∼95% probability of encountering a novel out-of-frame stop codon). On a per patient basis, an average of 0.002% of somatic variants overlapping transcripts were phased with a non-stop variant (range: 0-0.13%).

### Software performance and benchmarking

We attempted to compare the performance of `neoepiscope` with all other publicly available neoepitope prediction tools, but were unable to recapitulate several of these pipelines despite availability of the code. For example, the local version of CloudNeo had several Dockerfile and code-level errors that prevented us from running the installed tool. This raises a significant reproducibility and generalizability issue among many existing software tools. However, we were able to successfully benchmark four popular neoepitope prediction tools (pVACseq, and MuPeXI, NeoPredPipe, and TSNAD) along with `neoepiscope`.

Run as a full pipeline, `neoepiscope` outperformed pVACseq by an average of 44.7minutes per sample and outperformed MuPeXI by 3.4 minutes per sample, even when including germline variant calling (see Figure 5). NeoPredPipe and TSNAD outperformed `neoepiscope` by an average of 79.5, and 74.9 minutes per sample, respectively. These differences were predominantly accounted for by the average 76.7 minutes per sample runtime of germline variant calling that is required prior to running `neoepiscope`, as well as the additional 1.5 minutes per sample runtime of HapCUT2 when phasing somatic and germline variants. Excluding any pre-processing steps, `neoepiscope` outperformed pVACseq, MuPeXI, and TSNAD by 38.8, 4.86, and 3.22 minutes, respectively, and was outperformed by NeoPredPipe by 1.37 minutes.

We also compared the neoepitope sequences predicted by each tool. While the majority of neoepitopes predicted across all tools (55.0%) are consistent across all prediction tools, there are significant nuances and differences in predictions between tools (Figure 6). Of concern, we find that 22.9% of total neoepitopes are missed by TSNAD due to incomplete transcript/variant annotation, while 0.6% are erroneously reported due to incorrect variant interpretation. NeoPredPipe faces similar issues, with 4.8% of total neoepitopes missed due to incomplete annotation, and an additional 14.8% of neoepitopes resulting from incorrect variant interpretation. MuPeXI also reports 0.3% of neoepitopes due to incorrect variant interpretation. Among neoepitope prediction tools, only `neoepiscope`, MuPeXI, and NeoPredPipe are able to process peptide sequences resulting from a disrupted stop codon. For instance, pVACseq, TSNAD, and NeoPredPipe are unable to identify the 0.08% of total neoepitopes resulting from frameshift variants that extend the open reading frame past a stop codon. Similarly, pVACseq, TSNAD, and MuPeXI are unable to identify the 0.01% of neoepitopes resulting from stop codon point mutations.

Overall, 1% of all predicted neoepitopes result from phased variants and are predicted by both `neoepiscope` and pVACseq but not other tools. A similarly-sized set (0.8%) of neoepitopes resulting from phased variants is identified by `neoepiscope` alone. Of these, 7.9% appear to be due to inconsistencies in variant phasing between HapCUT2 and GATK ReadBackedPhasing; these neoepitopes are flagged as uncertain. However, the other 92.1% of these neoepitopes are phased consistently by both HapCUT2 and GATK ReadBackedPhasing, but are incorrectly missing from pVACseq output. Conversely, 3.3% of all neoepitopes are incorrectly reported (false positives), as they are predicted to be absent when taking variant phasing into account. Of these, 2.6% appear to be due to inconsistencies in variant phasing between HapCUT2 and GATK ReadBackedPhasing and are flagged as uncertain. However, the remaining 97.4% are consistently phased by both HapCUT2 and GATK ReadBackedPhasing, but erroneously reported by pVACseq and all other tools with the exception of `neoepiscope`. Taken together, a total of ∼5% of all neoepitopes are incorrectly or incompletely predicted without appropriately accounting for variant phasing.

There are several other anticipated discrepancies between the neoepitope prediction tools. For instance, there were nine neoepitopes derived from a mutation to a transcript from an unlocalized contig (chr1_KI270711v1_random) reported only by pVACseq and 36 associated with presumed RNA-editing events (on transcripts ENST00000361453 and ENST00000362079) which were reported only by pVACseq and TSNAD due to discrepancies in genomic annotation and variant effect prediction, respectively. However, in the absence of RNA-seq data, there is no way to reasonably differentiate performance for these sites. For the purposes of this analysis, we considered neoepitopes arising from nonsense-mediated decay, polymorphic pseudogene sequences, IG V and TR V transcripts to be bonafide sources of antigenic peptides [48], however `neoepiscope` can be parameterized to include or exclude any of these sequences as desired for a more nuanced interpretation of variant effects on putative neoepitopes.

## DISCUSSION

We report here a novel, performant, and flexible pipeline for neoepitope prediction from DNA-seq data (`neoepiscope`), which we demonstrate improves upon existing pipelines in several ways. In particular, we demonstrate the importance of germline context and variant phasing for accurate and comprehensive neoepitope identification. Not only do co-occurring somatic variants represent a source of previously unappreciated neoepitopes, but these compound epitopes may harbor more immunogenic potential compared to peptides from isolated somatic variants (due to multiple amino acid changes within the same epitope) [49,50]. Furthermore, we find that `neoepiscope` improves both sensitivity and specificity compared with existing software (which incorrectly or incompletely predicts ∼5% of neoepitopes [see Methods]). Indeed, this study is the first to critically evaluate and benchmark multiple neoepitope prediction tools, and raises the specter of broad reproducibility, accuracy, and usability issues within the field. Finally, we note that the `neoepiscope` framework is sufficiently flexible to accommodate numerous variant types, nonsense-mediated decay products, and epitope prediction across different genomes.

There are several limitations to `neoepiscope` at present. For instance, the software does not currently leverage RNA-seq data for gene expression or transcript-level variant phasing [51]. While the importance of gene dosing for neoepitope presentation remains unclear at this time, incorporation of transcript-level variant quantification could potentially improve accuracy of neoepitope prediction. Additionally, `neoepiscope` does not currently predict neoepitopes that may arise from alternative splicing events, RNA-editing, structural variation or gene fusions which are common across multiple cancer types [4,52–54]. Because `neoepiscope` implicitly assumes a diploid genome when using HapCUT2 for variant phasing, it may have decreased predictive accuracy in the context of significant somatic copy number alterations, though we do not observe this to be the case in the datasets presented herein [55]. Additionally, the software does not account for potentiation of the nonsense-mediated decay pathway by frameshifting indels, leading to uncertain implications for downstream neoepitope presentation in such cases. Lastly, as there remains significant uncertainty in predicting antigenicity *in vivo*, `neoepiscope` does not attempt to prioritize or rank putative neoepitopes in any way, relying instead on the end user to interpret results according to their own preferred schema.

In the future, we will address these limitations by incorporating downstream predictive methodologies which could aid in identifying the most immunogenic candidates (e.g., for personalized vaccine development [56,57]) and incorporating predictive features from RNA-seq as above to expand the repertoire of predicted neoepitopes. Given the proliferation of predictive tools, and their notable caveats and limitations as described in this study, we believe the field must advocate for the highest of standards in software reproducibility and transparency. Finally, we assert that future analyses should account for both germline context and variant phasing, and are happy to provide `neoepiscope` as a resource for the community, accordingly.

## AVAILABILITY

`neoepiscope` is available on GitHub at https://github.com/pdxgx/neoepiscope under the MIT license. Scripts for reproducing results described in the text are available at https://github.com/pdxgx/neoepiscope-paper under the MIT license.

## ACKNOWLEDGEMENTS

We would like to thank Mihir Paralkar and Mayur Paralkar for their support at `neoepiscope`’s inception. We would also like to thank Dr. Rachel Karchin for her group’s support with MHCnuggets integration. We finally thank Julianne David for her critical review of the manuscript.

## DISCLAIMER

The contents do not represent the views of the U.S. Department of Veterans Affairs or the United States government.

## SUPPLEMENTARY DATA

**Supplementary Table 1:**
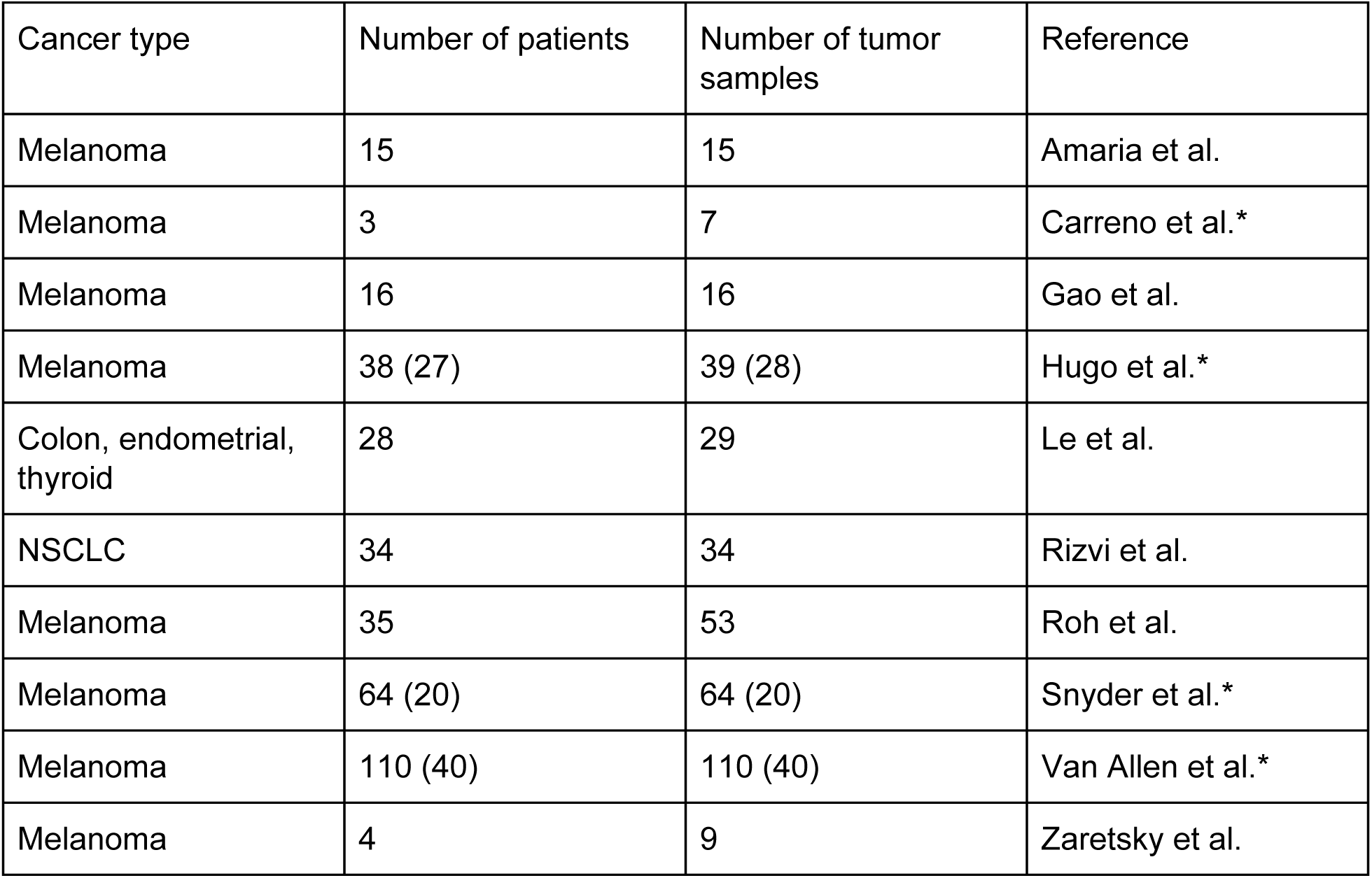
Summary of patients samples used to identify trends in variant phasing. Publicly available WES data from 10 studies was used to determine the frequency with which somatic variants are phased with germline or other somatic variants (see Materials and Methods). We summarize the study which produced each data set, the cancer types represented, and the number of patients/tumor samples sequenced. Studies that had complementary RNA sequencing reads available for at least a subset of patients are indicated by an asterisk in the “Reference” column, and the number of samples with complementary RNA sequencing data are indicated in parentheses in the “Number of patients” and “Number of tumor samples” columns if different than the number of samples with WES.

**Supplementary Figure 1.**
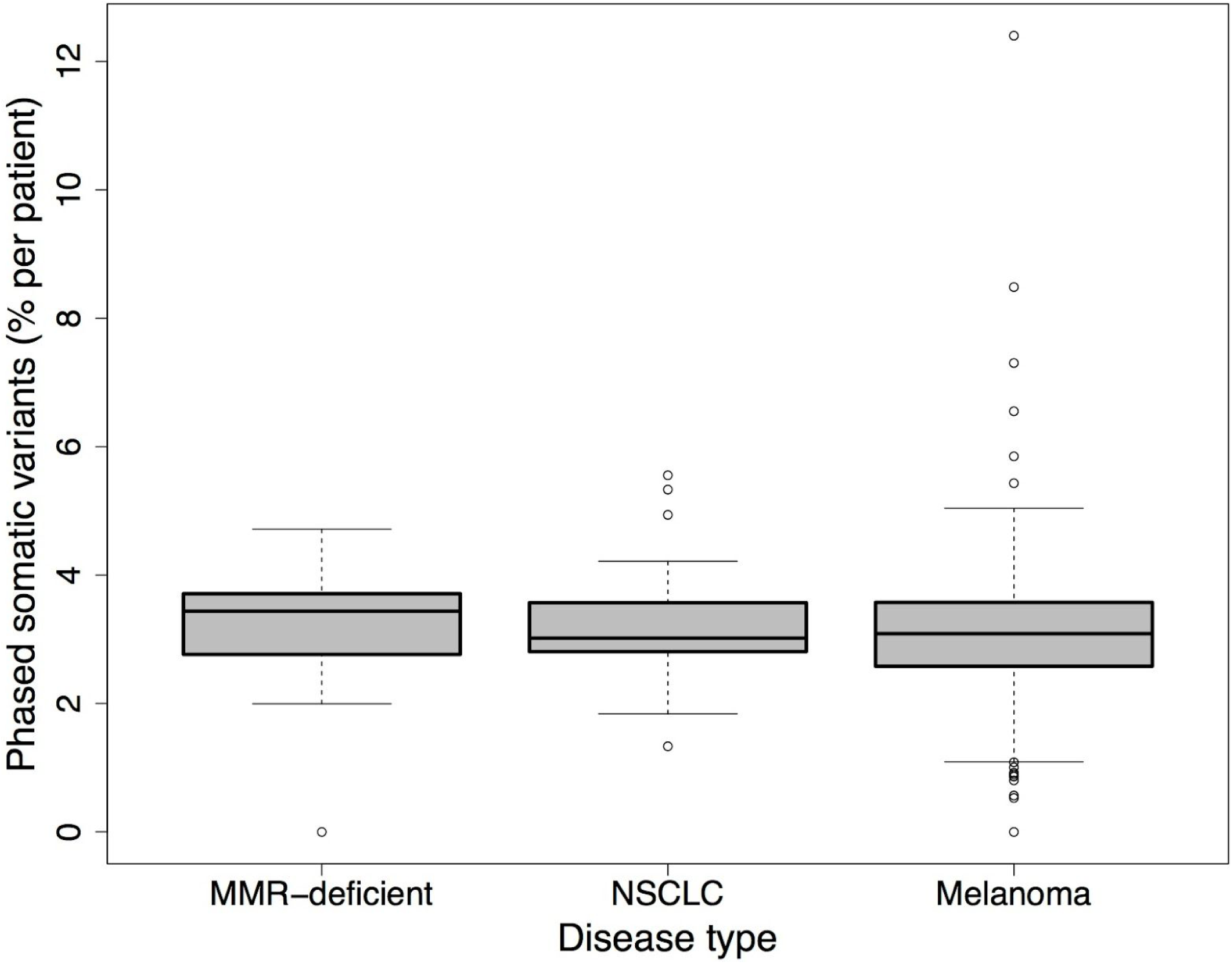
Co-occurrence of somatic variants by disease type. Box plots demonstrate the per-patient percentage of somatic variants (y-axis) across all tumors that co-occur with either germline variants or other somatic variants across 285 melanoma patients, 34 NSCLC patients, and 28 colon, endometrial, and thyroid cancer patients.

**Supplementary Figure 2.**
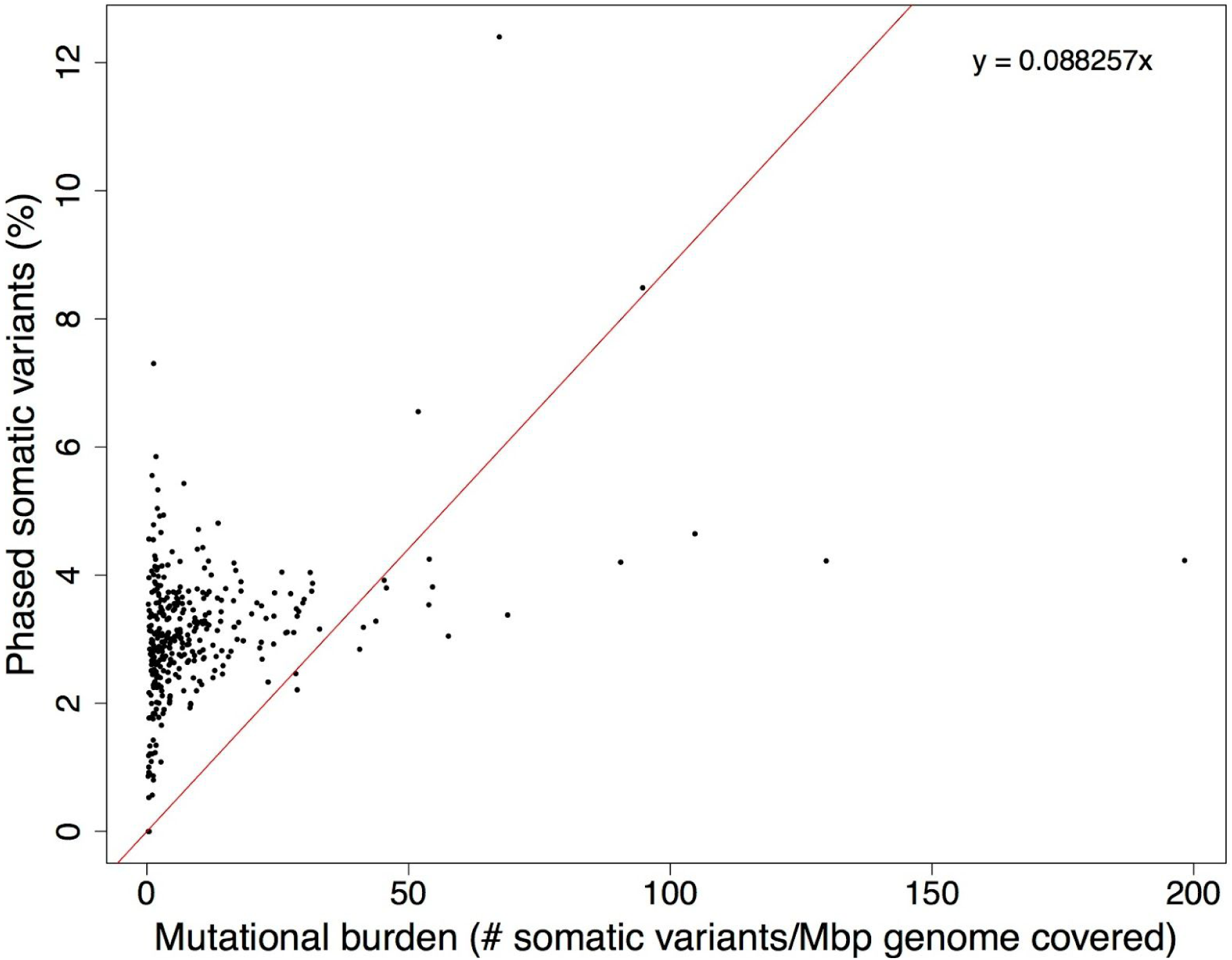
Relationship between mutational burden and phasing of somatic variants. The X-axis shows the coverage-adjusted mutation burden per patient (see Methods), while the Y-axis shows the percentage of somatic variants per patient that co-occur with a germline or another somatic variant within 72 bp. The best fit line through the origin is shown in red, with the equation describing it in the top right corner.

## REFERENCES

1. Efremova M, Finotello F, Rieder D, Trajanoski Z. Neoantigens Generated by Individual Mutations and Their Role in Cancer Immunity and Immunotherapy. Front Immunol. 2017;8: 1679.

2. Duan F, Duitama J, Al Seesi S, Ayres CM, Corcelli SA, Pawashe AP, et al. Genomic and bioinformatic profiling of mutational neoepitopes reveals new rules to predict anticancer immunogenicity. J Exp Med. 2014;211: 2231–2248.

3. Vogelstein B, Papadopoulos N, Velculescu VE, Zhou S, Diaz LA Jr, Kinzler KW. Cancer genome landscapes. Science. 2013;339: 1546–1558.

4. Zhang J, Mardis ER, Maher CA. INTEGRATE-neo: a pipeline for personalized gene fusion neoantigen discovery. Bioinformatics. 2017;33: 555–557.

5. Zhou Z, Lyu X, Wu J, Yang X, Wu S, Zhou J, et al. TSNAD: an integrated software for cancer somatic mutation and tumour-specific neoantigen detection. R Soc Open Sci. 2017;4: 170050.

6. Hundal J, Carreno BM, Petti AA, Linette GP, Griffith OL, Mardis ER, et al. pVAC-Seq: A genome-guided in silico approach to identifying tumor neoantigens. Genome Med. 2016;8: 11.

7. Bjerregaard A-M, Nielsen M, Hadrup SR, Szallasi Z, Eklund AC. MuPeXI: prediction of neo-epitopes from tumor sequencing data. Cancer Immunol Immunother. 2017;66: 1123–1130.

8. Turajlic S, Litchfield K, Xu H, Rosenthal R, McGranahan N, Reading JL, et al. Insertion-and-deletion-derived tumour-specific neoantigens and the immunogenic phenotype: a pan-cancer analysis. Lancet Oncol. 2017;18: 1009–1021.

9. 1000 Genomes Project Consortium, Abecasis GR, Altshuler D, Auton A, Brooks LD, Durbin RM, et al. A map of human genome variation from population-scale sequencing. Nature. 2010;467: 1061–1073.

10. Koire A, Kim YW, Wang J, Katsonis P, Jin H, Lichtarge O. Codon-level co-occurrences of germline variants and somatic mutations in cancer are rare but often lead to incorrect variant annotation and underestimated impact prediction. PLoS One. 2017;12: e0174766.

11. Muiño JM, Kuruoğlu EE, Arndt PF. Evidence of a cancer type-specific distribution for consecutive somatic mutation distances. Comput Biol Chem. 2014;53 Pt A: 79–83.

12. Szolek A, Schubert B, Mohr C, Sturm M, Feldhahn M, Kohlbacher O. OptiType: precision HLA typing from next-generation sequencing data. Bioinformatics. 2014;30: 3310–3316.

13. Li H, Durbin R. Fast and accurate long-read alignment with Burrows-Wheeler transform. Bioinformatics. 2010;26: 589–595.

14. Langmead B, Salzberg SL. Fast gapped-read alignment with Bowtie, 2. Nat Methods. 2012;9: 357–359.

15. Ewing AD, Houlahan KE, Hu Y, Ellrott K, Caloian C, Yamaguchi TN, et al. Combining tumor genome simulation with crowdsourcing to benchmark somatic single-nucleotide-variant detection. Nat Methods. 2015;12: 623–630.

16. McKenna A, Hanna M, Banks E, Sivachenko A, Cibulskis K, Kernytsky A, et al. The Genome Analysis Toolkit: a MapReduce framework for analyzing next-generation DNA sequencing data. Genome Res. 2010;20: 1297–1303.

17. Koboldt DC, Zhang Q, Larson DE, Shen D, McLellan MD, Lin L, et al. VarScan 2: somatic mutation and copy number alteration discovery in cancer by exome sequencing. Genome Res. 2012;22: 568–576.

18. Edge P, Bafna V, Bansal V. HapCUT2: robust and accurate haplotype assembly for diverse sequencing technologies. Genome Res. 2017;27: 801–812.

19. Langmead B, Trapnell C, Pop M, Salzberg SL. Ultrafast and memory-efficient alignment of short DNA sequences to the human genome. Genome Biol. 2009;10: R25.

20. O’Donnell TJ, Rubinsteyn A, Bonsack M, Riemer AB, Laserson U, Hammerbacher J. MHCflurry: Open-Source Class I MHC Binding Affinity Prediction. Cell Syst. 2018;7: 129–132.e4.

21. Bhattacharya R, Sivakumar A, Tokheim C, Guthrie VB, Anagnostou V, Velculescu VE, et al. Evaluation of machine learning methods to predict peptide binding to MHC Class I proteins [Internet]., 2017. doi:10.1101/154757

22. Nielsen M, Lundegaard C, Blicher T, Peters B, Sette A, Justesen S, et al. Quantitative Predictions of Peptide Binding to Any HLA-DR Molecule of Known Sequence: NetMHCIIpan. PLoS Comput Biol. 2008;4: e1000107.

23. Zaretsky JM, Garcia-Diaz A, Shin DS, Escuin-Ordinas H, Hugo W, Hu-Lieskovan S, et al. Mutations Associated with Acquired Resistance to PD-1 Blockade in Melanoma. N Engl J Med. 2016;375: 819–829.

24. Carreno BM, Magrini V, Becker-Hapak M, Kaabinejadian S, Hundal J, Petti AA, et al. A dendritic cell vaccine increases the breadth and diversity of melanoma neoantigen-specific T cells. Science. 2015;348: 803–808.

25. Hugo W, Zaretsky JM, Sun L, Song C, Moreno BH, Hu-Lieskovan S, et al. Genomic and Transcriptomic Features of Response to Anti-PD-1 Therapy in Metastatic Melanoma. Cell. 2017;168: 542.

26. Snyder A, Makarov V, Merghoub T, Yuan J, Zaretsky JM, Desrichard A, et al. Genetic basis for clinical response to CTLA-4 blockade in melanoma. N Engl J Med. 2014;371: 2189–2199.

27. Van Allen EM, Miao D, Schilling B, Shukla SA, Blank C, Zimmer L, et al. Genomic correlates of response to CTLA-4 blockade in metastatic melanoma. Science. 2015;350: 207–211.

28. Gao J, Shi LZ, Zhao H, Chen J, Xiong L, He Q, et al. Loss of IFN-γ Pathway Genes in Tumor Cells as *a Mechanism of Resistance to Anti-CTLA*-4 Therapy. Cell. 2016;167: 397–404.e9.

29. Roh W, Chen P-L, Reuben A, Spencer CN, Prieto PA, Miller JP, et al. Integrated molecular analysis of tumor biopsies on sequential CTLA-4 and PD-1 blockade reveals markers of response and resistance. Sci Transl Med. 2017;9.doi:10.1126/scitranslmed.aah3560

30. Amaria RN, Reddy SM, Tawbi HA, Davies MA, Ross MI, Glitza IC, et al. Neoadjuvant immune checkpoint blockade in high-risk resectable melanoma. Nat Med. 2018;24: 1649–1654.

31. Rizvi NA, Hellmann MD, Snyder A, Kvistborg P, Makarov V, Havel JJ, et al. Cancer immunology. Mutational landscape determines sensitivity to PD-1 blockade in non-small cell lung cancer. Science. 2015;348: 124–128.

32. Le DT, Durham JN, Smith KN, Wang H, Bartlett BR, Aulakh LK, et al. Mismatch repair deficiency predicts response of solid tumors to PD-1 blockade. Science. 2017;357: 409–413.

33. cancerit. cancerit/dockstore-cgpmap. In: GitHub [Internet]. [cited 12 Sep 2018]. Available: https://github.com/cancerit/dockstore-cgpmap

34. Li H, Durbin R. Fast and accurate short read alignment with Burrows-Wheeler transform. Bioinformatics. 2009;25: 1754–1760.

35. gt. gt1/biobambam2. In: GitHub [Internet]. [cited 12 Sep 2018]. Available: https://github.com/gt1/biobambam2

36. Fan Y, Xi L, Hughes DST, Zhang J, Zhang J, Futreal PA, et al. MuSE: accounting for tumor heterogeneity using a sample-specific error model improves sensitivity and specificity in mutation calling from sequencing data. Genome Biol. 2016;17: 178.

37. Cibulskis K, Lawrence MS, Carter SL, Sivachenko A, Jaffe D, Sougnez C, et al. Sensitive detection of somatic point mutations in impure and heterogeneous cancer samples. Nat Biotechnol. 2013;31: 213–219.

38. Ye K, Schulz MH, Long Q, Apweiler R, Ning Z. Pindel: a pattern growth approach to detect break points of large deletions and medium sized insertions from paired-end short reads. Bioinformatics. 2009;25: 2865–2871.

39. Radenbaugh AJ, Ma S, Ewing A, Stuart JM, Collisson EA, Zhu J, et al. RADIA: RNA and DNA integrated analysis for somatic mutation detection. PLoS One. 2014;9: e111516.

40. Larson DE, Harris CC, Chen K, Koboldt DC, Abbott TE, Dooling DJ, et al. SomaticSniper: identification of somatic point mutations in whole genome sequencing data. Bioinformatics. 2012;28: 311–317.

41. Ellrott K, Bailey MH, Saksena G, Covington KR, Kandoth C, Stewart C, et al. Scalable Open Science Approach for Mutation Calling of Tumor Exomes Using Multiple Genomic Pipelines. Cell Syst. 2018;6: 271–281.e7.

42. Tan A, Abecasis GR, Kang HM. Unified representation of genetic variants. Bioinformatics. 2015;31: 2202–2204.

43. Quinlan AR, Hall IM. BEDTools: a flexible suite of utilities for comparing genomic features. Bioinformatics. 2010;26: 841–842.

44. Dobin A, Davis CA, Schlesinger F, Drenkow J, Zaleski C, Jha S, et al. STAR: ultrafast universal RNA-seq aligner. Bioinformatics. 2013;29: 15–21.

45. Bassani-Sternberg M, Bräunlein E, Klar R, Engleitner T, Sinitcyn P, Audehm S, et al. Direct identification of clinically relevant neoepitopes presented on native human melanoma tissue by mass spectrometry. Nat Commun. 2016;7: 13404.

46. McLaren W, Gil L, Hunt SE, Riat HS, Ritchie GRS, Thormann A, et al. The Ensembl Variant Effect Predictor. Genome Biol. 2016;17: 122.

47. Nielsen M, Lundegaard C, Blicher T, Lamberth K, Harndahl M, Justesen S, et al. NetMHCpan, a Method for Quantitative Predictions of Peptide Binding to Any HLA-A and -B Locus Protein of Known [Internet]. SciVee., 2007. doi:10.4016/4651.01

48. Apcher S, Daskalogianni C, Lejeune F, Manoury B, Imhoos G, Heslop L, et al. Major source of antigenic peptides for the MHC class I pathway is produced during the pioneer round of mRNA translation. Proc Natl Acad Sci U S A. 2011;108: 11572–11577.

49. Pradeu T, Carosella ED. On the definition of a criterion of immunogenicity. Proc Natl Acad Sci U S A. 2006;103: 17858–17861.

50. Yadav M, Jhunjhunwala S, Phung QT, Lupardus P, Tanguay J, Bumbaca S, et al. Predicting immunogenic tumour mutations by combining mass spectrometry and exome sequencing. Nature. 2014;515: 572–576.

51. Castel SE, Mohammadi P, Chung WK, Shen Y, Lappalainen T. Rare variant phasing and haplotypic expression from RNA sequencing with phASER. Nat Commun. 2016;7: 12817.

52. Smart AC, Margolis CA, Pimentel H, He MX, Miao D, Adeegbe D, et al. Intron retention as a novel source of cancer neoantigens [Internet]., 2018. doi:10.1101/309450

53. Kahles A, Lehmann K-V, Toussaint NC, Hüser M, Stark SG, Sachsenberg T, et al. Comprehensive Analysis of Alternative Splicing Across Tumors from 8,705 Patients. Cancer Cell. 2018;34: 211–224.e6.

54. Xu X, Wang Y, Liang H. The role of A-to-I RNA editing in cancer development. Curr Opin Genet Dev. 2018;48: 51–56.

55. Zack TI, Schumacher SE, Carter SL, Cherniack AD, Saksena G, Tabak B, et al. Pan-cancer patterns of somatic copy number alteration. Nat Genet. 2013;45: 1134–1140.

56. Ott PA, Hu Z, Keskin DB, Shukla SA, Sun J, Bozym DJ, et al. An immunogenic personal neoantigen vaccine for patients with melanoma. Nature. 2017;547: 217–221.

57. Sahin U, Derhovanessian E, Miller M, Kloke B-P, Simon P, Löwer M, et al. Personalized RNA mutanome vaccines mobilize poly-specific therapeutic immunity against cancer. Nature. 2017;547: 222–226.

58. Bais P, Namburi S, Gatti DM, Zhang X, Chuang JH. CloudNeo: a cloud pipeline for identifying patient-specific tumor neoantigens. Bioinformatics. 2017;33: 3110–3112.

59. Tang S, Madhavan S. neoantigenR: An annotation based pipeline for tumor neoantigen identification from sequencing data [Internet]., 2017. doi:10.1101/171843

60. Chang T-C, Carter RA, Li Y, Li Y, Wang H, Edmonson MN, et al. The neoepitope landscape in pediatric cancers. Genome Med. 2017;9: 78.

61. Rubinsteyn A, Kodysh J, Hodes I, Mondet S, Aksoy BA, Finnigan JP, et al. Computational Pipeline for the PGV-001 Neoantigen Vaccine Trial. Front Immunol. 2017;8: 1807.

62. Rech AJ, Balli D, Mantero A, Ishwaran H, Nathanson KL, Stanger BZ, et al. Tumor Immunity and Survival as a Function of Alternative Neopeptides in Human Cancer. Cancer Immunology Research. 2018;6: 276–287.

63. Schenck RO, Lakatos E, Gatenbee C, Graham TA, Anderson ARA. NeoPredPipe: High-Throughput Neoantigen Prediction and Recognition Potential Pipeline [Internet]., 2018. doi:10.1101/409839

64. Kim S, Kim HS, Kim E, Lee MG, Shin E-C, Paik S, et al. Neopepsee: accurate genome-level prediction of neoantigens by harnessing sequence and amino acid immunogenicity information. Ann Oncol. 2018;29: 1030–1036.

65. Hundal J, Kiwala S, Feng Y-Y, Liu CJ, Govindan R, Chapman WC, et al. Accounting for proximal variants improves neoantigen prediction. Nat Genet. 2019;51: 175–179.

